# A Mean-Field Description of Bursting Dynamics in Spiking Neural Networks with Short-Term Adaptation

**DOI:** 10.1101/806273

**Authors:** R. Gast, H. Schmidt, T.R. Knösche

## Abstract

Bursting plays an important role in neural communication. At the population level, macroscopic bursting has been identified in populations of neurons that do not express intrinsic bursting mechanisms. For the analysis of such phase transitions, mean-field descriptions of macroscopic bursting behavior pose a valuable tool. In this article, we derive mean-field descriptions of populations of spiking neurons in which collective bursting behavior arises via short-term adaptation mechanisms. Specifically, we consider synaptic depression and spike-frequency adaptation in networks of quadratic integrate-and-fire neurons. We characterize the emerging bursting behavior using bifurcation analysis and validate our mean-field derivations by comparing the microscopic and macroscopic descriptions of the population dynamics. Hence, we provide mechanistic descriptions of phase transitions between bursting and non-bursting population dynamics which play important roles in both healthy neural communication and neurological disorders.

## I. INTRODUCTION

The brain, composed of billions of single cells, has been demonstrated to possess a hierarchical, modular organization, indicative of a complex dynamical system^1^. Within this hierarchy, populations of neurons form functional entities, the states of which are defined by the collective dynamics of the population rather than by the activity of each single cell. Mean-field descriptions of the macroscopic dynamics of such populations are a valuable tool for the mathematical analysis of collective phenomena as well as for computational models of multiple coupled populations of neurons. Population bursting is a particular mode of collective behavior that plays a major role in both healthy and pathological neural dynamics. In healthy neural communication bursting activity may allow for a more reliable information transmission via chemical synapses^2^. This can be explained by the synchronized activity of the population during the burst, which stabilizes neural information transmission against different types of noise^3^. On the other hand, increased bursting activity has been found in various neurological diseases such as epilepsy or Parkinson’s disease and may thus also act disruptively on neural communication, if exceeding certain levels of occurrence^4,5^. Thus, mean-field models of collective bursting are important for theoretical investigations of information transmission between neural populations as well as transitions between healthy and pathological states of population dynamics. The aim of this article is to provide and validate mean-field descriptions for collective bursting emerging from the dynamic interaction of short-term adaptation mechanisms and recurrent excitation in populations of coupled spiking neurons.

At the single neuron level, bursting is characterized by the neuron firing a group of subsequent spikes, followed by a period of quiescence^6^. This behavior has been suggested to result from adaptive mechanisms introducing a slow time scale which enables dynamic regimes of bursting and controls the burst period^6,7^. Mathematical descriptions of such adaptation mechanisms have been developed accordingly at the level of single cells. Importantly, bursting has also been reported in populations of cells without intrinsic bursting mechanisms^6,8,9^. In such cases, bursting can be conceived as a property of the collective dynamic interactions within the population. The mechanisms behind emergent bursting are not well understood, however, since most of the computational literature on bursting focuses on single cells^6,10^. Among the few studies on emergent bursting, a common approach to model bursting at the population level is to globally couple an excitatory with an inhibitory population^9,11^. Other suggested bursting mechanisms include the action of neuromodulators^8^, feed-forward inhibition^12^ and spike-frequency adaptation (SFA)^13^. For example, Van Vreeswijk et al. demonstrated in a network of coupled, excitatory leaky integrate-and-fire neurons that SFA can lead to the emergence of network bursting^13^. Typically, such investigations employ numerical analyses of single cell networks, where the macroscopic state variables have to be inferred from the single cell activities. However, a direct mathematical description of the macroscopic dynamics would be beneficial for both mathematical analyses of emergent bursting and studies on multiple coupled bursting populations. Gigante and colleagues derived a mean-field description for the special case of SFA in a network of coupled linear integrate- and-fire neurons, e mploying t he F okker-Planck formalism and an adiabatic approximation given long SFA time scales^14^. Analyzing this mean-field description, they were able to identify two different types of collective bursting. In the present study, we generalize these findings and show under which conditions bursting can emerge as a collective phenomenon from mere short-term adaptation within an excitatory population of globally coupled quadratic integrate-and-fire (QIF) neurons. To this end, we employ a recently derived mean-field description of the macroscopic dynamics of such a QIF population^15^. We consider different short-term adaptation mechanisms that have been reported to affect neural excitability and incorporate them in the microscopic description of the QIF population dynamics. We show for which of these adaptation mechanisms a mean-field description can be derived according to the approach of Montbrió *et al.*^15^. Employing those mean-field descriptions, we analyze the emergence of states of collective bursting and identify boundary conditions for such bursting to occur.

## II. MODEL DEFINITION

The QIF neuron is the canonical form of type-I neurons and has previously been used in combination with linear adaptation as a basis for models of bursting cells^6^. The evolution equation of the membrane potential *V*_*i*_ of a single QIF neuron *i* is given by

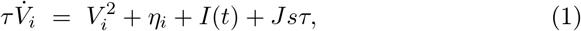

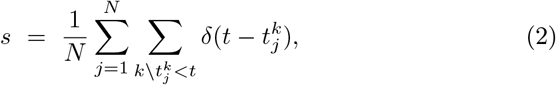

with background current *η*_*i*_, synaptic strength *J*, evolution time constant *τ*, extrinsic input *I*(*t*) and synaptic activation *s*. The latter represents instantaneous synaptic coupling in an all-to-all coupled network of N neurons. A neuron *i* emits its *k*^*th*^ spike at time 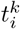 when it reaches *V*_*θ*_ upon which *V*_*i*_ is reset to *V*_*r*_. Recently, Montbrió *et al.*^15^ have shown that there exists an exact mean-field description for the macroscopic dynamics of the population dynamics given by (1) and (2) in the limit of *N* → ∞ and *V*_*θ*_ = −*V*_*r*_ → ∞. These can be expressed in terms of the evolution of the average firing rate *r* and membrane potential *v*.

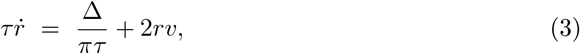

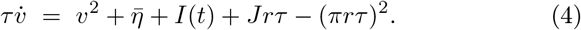

Here, 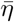 and Δ are the center and full width at half maximum of a Lorentzian distribution over the single neuron parameters *η*_*i*_. Furthermore, synaptic conversion is assumed to be instantaneous, which allows to set *r* = *s*. For simplicity, we set the evolution time constant to *τ* = 1 and define all other time-dependent variables used throughout this article in units of *τ*. As shown in^15^, the variables *V* of the microscopic system always follow a Lorentzian distribution,

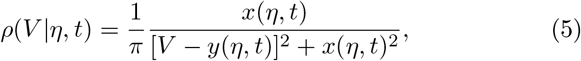

where *x*(*η*, *t*) is associated with the firing rate *r*(*η*, *t*) via *x*(*η*, *t*) = *πr*(*η*, *t*), and *y*(*η*, *t*) is the mean of the membrane potential *v*(*η*, *t*). Note that *x*(*η*, *t*) and *y*(*η*, *t*) are microscopic variables pertaining to neurons with specific *η*. The microscopic system follows the continuity equation

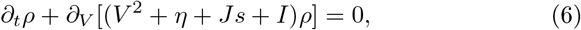

and by inserting eq. 5 into eq. 6 one obtains the dynamics of *x*(*η*, *t*) and *y*(*η*, *t*) in complex form

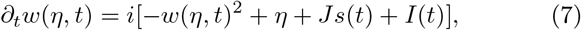

with *w*(*η*, *t*) = *x*(*η*, *t*)+*iy*(*η*, *t*). Eq. 7 can then be reduced to 3 using the Lorentzian distribution

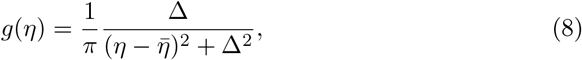

given in complex form by

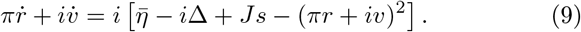

As shown by Montbrió *et al.*^15^, there exists a regime in which two stable states of the system defined by (3) and (4) can co-exist, a stable fixed point representing low activity and a stable focus representing high activity. To introduce bursting to this system, a mechanism is required that alternatingly switches between those two states.

## III. SYNAPTIC DEPRESSION

The first short-term adaptation mechanism we consider is synaptic depression (SD) which is a multiplicative down-scaling of the synaptic efficacy. Neurobiologically, this can be implemented in various ways, such as postsynaptic receptor desensitization, alterations in the density of post-synaptic receptors or resource depletion at the synapse^16,17^.

### III.A. Mathematical Definition of SD

To introduce SD to our system, we change (2) as follows

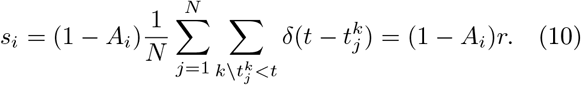

This adds a dependency of the post-synaptic activation *s*_*i*_ on an adaptation variable *A*_*i*_ pertaining to the *i*^*th*^ post-synaptic neuron. The latter follows the microscopic evolution equations

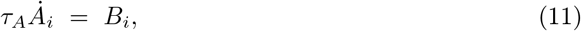

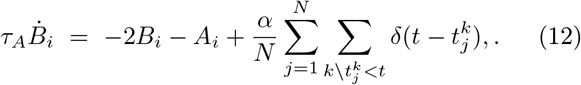

with adaptation rate *α* and time scale *τ*_*A*_. The adaptation dynamics given by (11) and (12) can be re-written as a convolution of the average firing rate *r* with an alpha kernel:

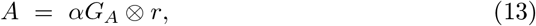

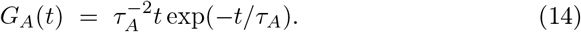

This choice accounts for the delay between post-synaptic activation and peak adaptation as well as for the slow decay of the adaptation to baseline that have been reported in experimental studies^16–18^. Importantly, this kind of adaptive mechanism has no effect on an isolated neuron, because it only affects the susceptibility to synaptic input which is effectively zero shortly after it spiked (due to refractoriness). Therefore, any changes to the network behavior caused by SD have to emerge from the interaction dynamics of the network units. Furthermore, the adaptation is coupled to the mean-field firing rate of the population, i.e. each spike in the network triggers post-synaptic adaptation at all network units. Hence, to introduce SD at the macroscopic level, we merely need to change (3) and (4) to

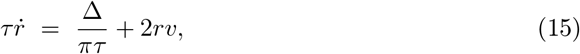

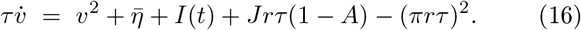

### III.B. Effects of SD

Next, we applied bifurcation analysis to the 4 dimensional system defined by (13–16). For different values of *α*, we initialized the model at a low activity state and continued the model in 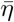. To this end, and for all other parameter continuations reported in this article, we used the software package AUTO-07p^19^. The adaptation time scale was chosen as *τ*_*A*_ = 10*τ*, corresponding to slow adaptation relative to the evolution of the average membrane potential and firing rate. In accordance with the analysis of Montbrió *et al.*^15^, we found two fold bifurcations for *α* = 0, defining the borders of a bi-stable regime in 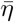. For an increasing adaptation rate *α*, we identified a parameter regime in which a supercritical and a subcritical Andronov-Hopf bifurcation occur (see Figure 1A). As shown in Figure 1B, the unstable limit cycle emerging from the subcritical Andronov-Hopf bifurcation turned into a stable limit cycle via a fold bifurcation. Further continuation of the stable region of the limit cycle in *η* revealed a boundary at which the limit cycle period grew towards infinity, indicative of a homoclinic bifurcation. The stable regime of the bursting limit cycle is visualized in green in Figure 1B. It can co-exist with the high-activity focus and hence allows for various transitions between bursting and steady-state behavior. As illustrated in Figure 1C-D, the stable bursting state can be transiently entered from either a low-activity state through excitation (Figure 1C) or from a high-activity state through inhibition (Figure 1D). Furthermore, the bi-stable regime allows for hysteresis, i.e. switching between limit cycle and focus equilibrium through transient excitatory and inhibitory inputs (Figure 1E). In neural communication, this regime is particularly relevant, since it allows for quick transitions between highly different firing modes via transient inputs and introduces a form of network memory. However, it is also of interest for pathological neural dynamics such as observed in epilepsy, which have been proposed to reflect switching between a healthy state of low neural synchrony and a co-existing pathological, synchronous state^20,21^. Importantly, Figure 1C-E show a close correspondence between numerical simulations of the macroscopic and the microscopic population descriptions.

**Fig. 1.**
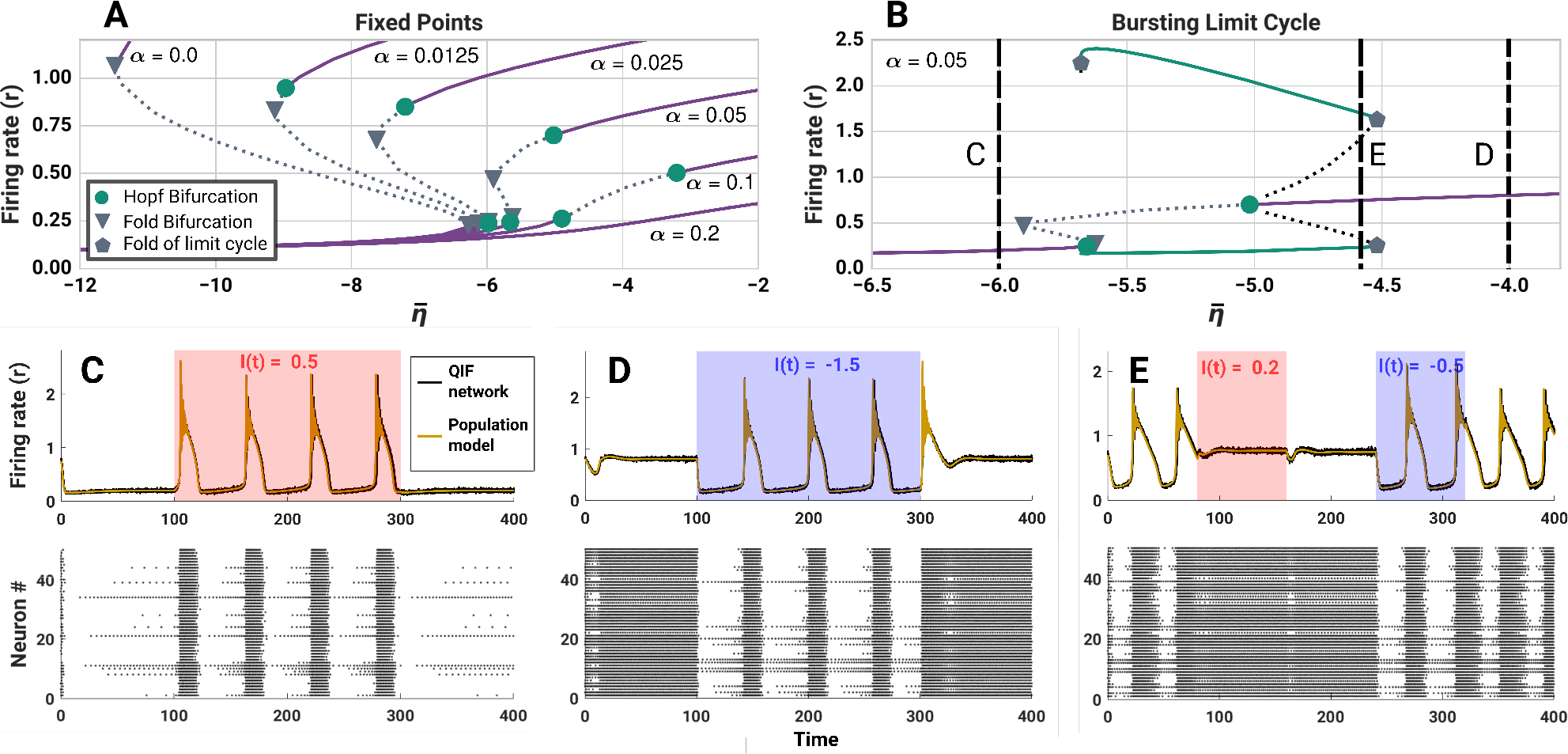
Bursting due to synaptic depression. **(A)** Bifurcation diagram of fixed points in *η* for various values of *α*. Stable fixed points are marked by solid lines whereas unstable fixed points are marked by dashed lines. **(B)** The subcritical Hopf bifurctions give rise to limit cycles representing bursting behavior. Minimum and maximum firing firing rates of the limit cycle are depicted in green. Fold bifurcations of the limit cycles delimit the bistable regime of bursting and regular firing. Vertical lines with letters indicate initialization points used for C-E, respectively. **(C)** Below the bistable regime, a positive stimulus leads to transient bursting. **(D)** Above the bistable regime, a negative stimulus leads to temporary bursting. **(E)** In the bistable regime, positive and negative stimuli switch the system between sustained bursting and sustained regular firing. Model parameters were set to Δ = 2, 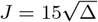, *τ* = 1, *τ*_*A*_ = 10.

### III.C. Limit cycle characteristics

For a better understanding of the bi-stable regime, we mapped out the basins of attraction with respect to all four state variables of the model. Figure 2 visualizes different trajectories of the system when initialized at different points near the unstable limit cycle that separates the limit cycle (bursting activity) and the stable fixed point (steady state activity). This unstable limit cycle behaves like a separatrix along the fast sections of its orbit. In fact, it can be understood as the cross-section through the actual hyperplane delimiting the two basins of attraction. However, near the slowly varying section of the trajectory, the unstable limit cycle maps out points that behave like a stable focus in the fast system.

**Fig. 2.**
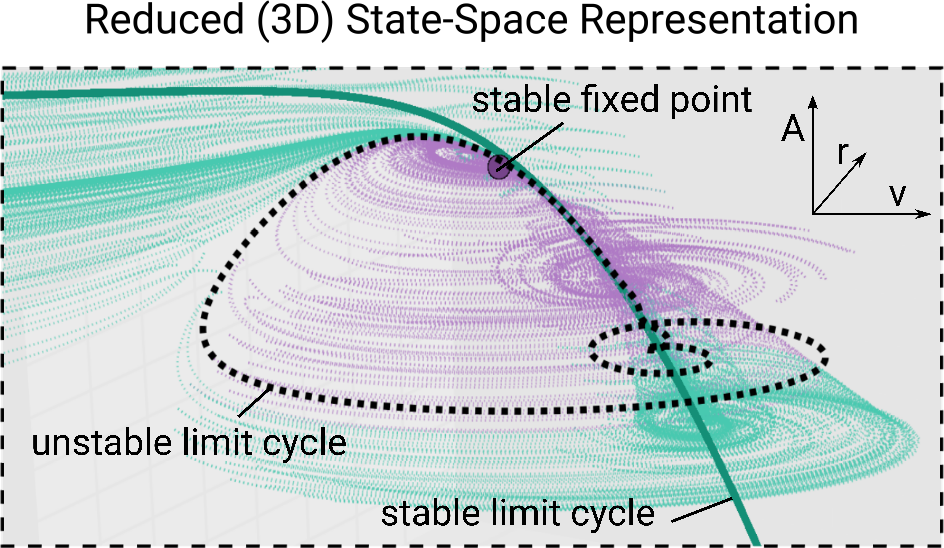
Co-existence of bursting and steady-state behavior. Reduced, 3-dimensional state-space representation of the system dynamics in *r*, *v*, and *A*. For the present parameters the stable limit cycle (bold green curve, representing bursting behavior) coexists with a stable focus (purple dot) and an unstable limit cycle (black dashed curve). Thin curves mark trajectories that end in the basin of attraction of the limit cycle (green) or the focus (purple). Model parameters: Δ = 2, 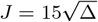, *τ* = 1.0, *τ*_*A*_ = 10 *α* = 0.05, 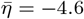.

In a next step, we performed a two-parameter continuation of the subcritical Andronov-Hopf bifurcation in 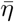 and *α*, to examine the dependence of population bursting on the interplay between network excitation and adaptation rate. From Figure 3A, we can conclude that dynamic regimes of stable bursting (marked in green) can be found for a substantial but limited range of the two parameters. This range is bounded by fold of limit cycle bifurcations that mark the disappearance of the stable limit cycle. For 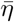, the parameter range in which the limit cycle exists corresponds to most of the cells in the population being in an excitable regime and has been reported for a number of models using QIF neurons (e.g.^15,22,23^). Within this range, the inter-burst period can be varied from 13*τ* up to 105*τ* via changes in *α* and 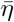 (see Figure 3B). Thus, the burst frequency scales with the input strength, which is a desirable property for encoding information about the population input via its activity. In summary, we identified SD as a potential mechanism for bursting to occur in networks of globally coupled spiking neurons. We demonstrated that this bursting mechanism could be transiently switched on and off via transient input currents and found that the inter-burst frequency can encode information about the input strength.

**Fig. 3.**
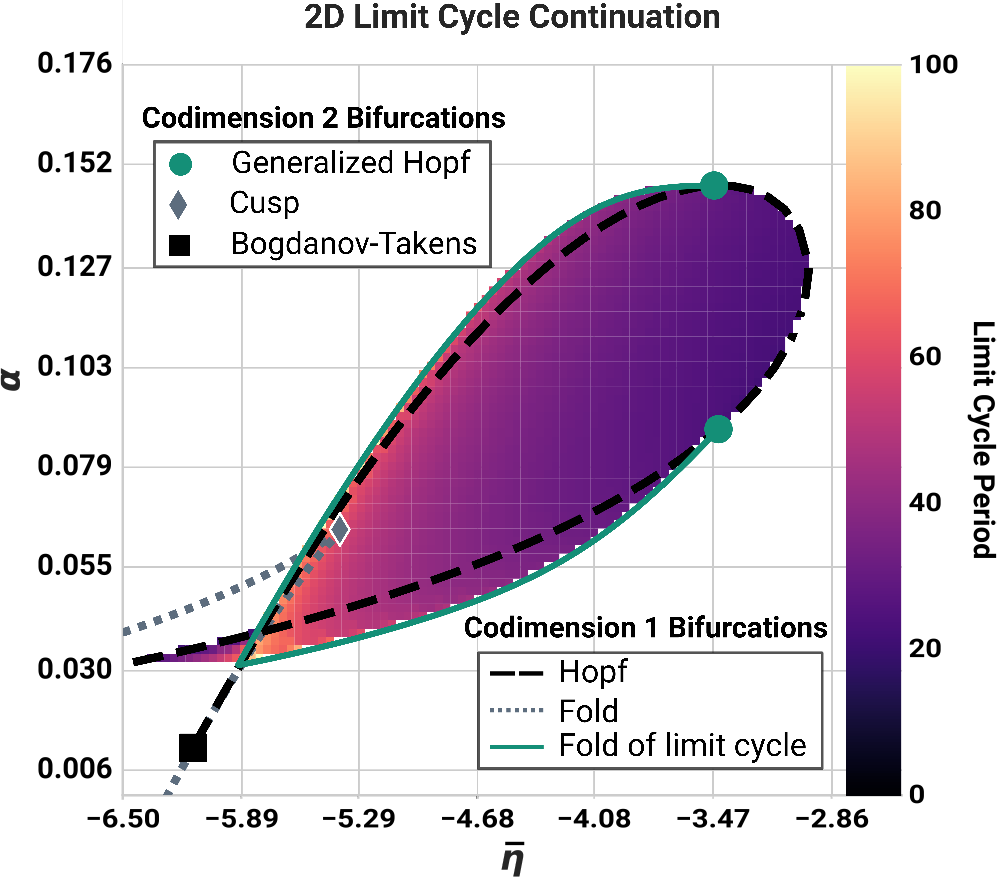
Existence and period of bursting. Lines indicate twoparameter continuations of codimension 1 bifurcations in the (*η*,*α*) plane. The color-coded region shows the inter-burst period of the stable limit cycle (depicted in units of *τ*).

### III.D. Pre-Synaptic SD

Since the mean-field description for SD has been derived under the assumption of post-synaptic adaptation, it cannot describe pre-synaptic multiplicative adaptation such as vesicle depletion^24^. To derive a mean-field description for such a case, a pre-synaptic adaptation variable *A*_*j*_ has to be considered:

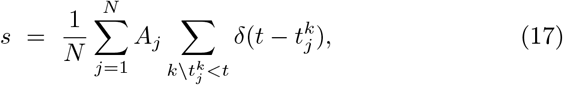

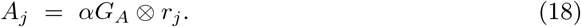

One may (naively) assume that *ρ*(*V*) still obeys the Lorentzian distribution eq. (5). Then, the dynamics of *w* would be given by

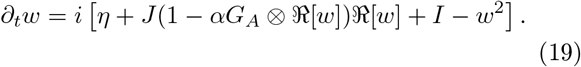

Here we make use of *s*_*i*_ = *x*(*η*_*i*_, *t*)/*π* again. This raises the problem of evaluating the integral containing the product of the microscopic firing rate with its temporal convolution. Also, with the firing rates being weighted in such a manner it is uncertain whether eq. (5) still holds, even if *τ*_*A*_ ≫ *τ*. Thus, it is not obvious how to derive the mean field description for this particular scenario.

## IV. SPIKE-FREQUENCY ADAPTATION

In this section, we examine how well our results for multiplicative adaptation generalize to additive shortterm adaptation mechanisms, known as spike-frequency adaptation (SFA). SFA differs from the above described adaptation mechanism in two aspects: (1) It affects the pre-synaptic activity instead of the post-synaptic efficacy, and (2) it acts additive instead of multiplicative^13,14^.

### IV.A. Mathematical Defition of SFA

SFA is a homeostatic mechanism that acts at the single cell level via spike-triggered balancing currents^10,25^. As such, SFA is an adaptive mechanism driven by the firing rate of a single cell rather than the firing rate of the whole network. Therefore, we introduce neuron-specific adaptation variables *A*_*i*_ and *B*_*i*_:

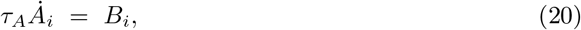

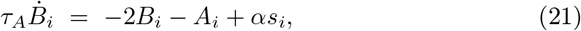

with a neuron-specific firing rate *s*_*i*_ given by

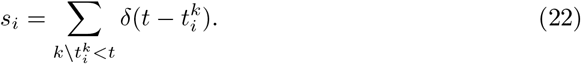

Adding the adaptation variable *A*_*i*_ to (1), we get the following evolution equation for the membrane potential of the single neuron:

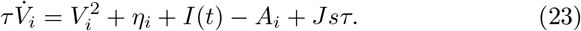

If *τ*_*A*_ ≫ *τ*, then we may assume that the adaption variable *A_i_* changes very slowly in comparison to *V*_*i*_, and that the Lorentzian ansatz (eq. (5)) holds. Effectively, the variable *A_i_* can be regarded as constant and be absorbed into *η*_*i*_ in this limit (for a similar approach, see^14^). We note here that the Lorentzian ansatz eq (5) is independent of the distribution of {*η*_*i*_}. The neuron-specific firing rate *s*_*i*_ is associated with *x*(*η*, *t*) via *s*_*i*_ = *x*(*η*_*i*_, *t*)/*π*. Hence, the microscopic dynamics are given by

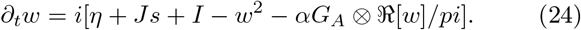

If *g*(*η*) follows the Lorentzian distribution, then eq. (24) results in

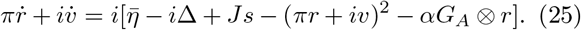

Thus, the pre-synaptic additive model is identical to a post-synaptic additive model at the macroscopic scale if *τ*_*A*_ ≫ *τ*. At the macroscopic scale, post-synaptic SFA is straightforward to realize by adding *A* on the r.h.s. of (3), leading to the macroscopic evolution equations

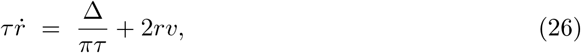

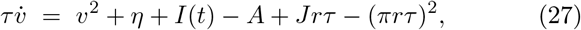

where *s* = *r* and *A* is still given by (13) and (14). Note that (26) and (27) are equivalent to the complex differential form given by (25).

### IV.B. Effects of SFA

Using the system defined by (13), (14), (26) and (27), we repeated the parameter continuation in 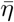 for different values of *α*. This was done to examine whether the results we obtained for SD-induced population bursting would translate to an additive adaptation mechanism. As can be seen in Figure 4A, we found results strikingly similar to the ones we found for SD. For sufficiently strong levels of SFA (parametrized via *α*), we found a subcritical Andronov-Hopf bifurcation marking the birth of a bursting limit cycle. Furthermore, we again found a bi-stable region, in which the bursting limit cycle co-exists with the stable focus, separated by an unstable limit cycle. These regimes could be well traversed via transient inputs, as depicted in Figure 4B. Driving the microscopic model given by (2) and (20–23) with the same transient inputs, we found that the spiking dynamics were still attracted to the low-dimensional manifold described by the macroscopic system. This shows, that even with *τ_A_* = 10_*τ*_, the condition *τ*_*A*_ ≫ *τ* is sufficiently satisfied for the macroscopic description of the population bursting dynamics to be valid.

**FIG. 4.**
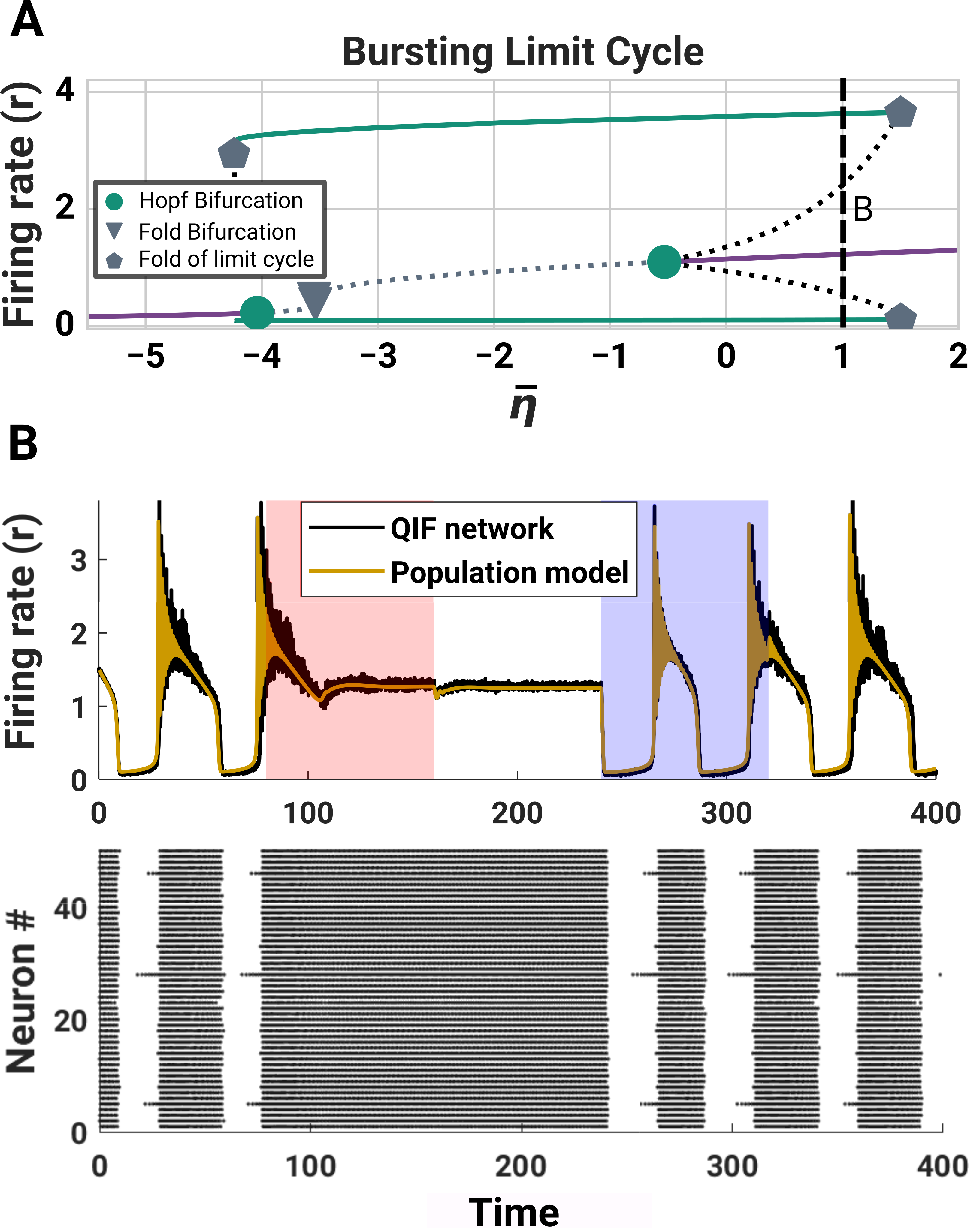
Bursting due to SFA. **(A)** Similar to SD, SFA changes fixedpoint structure and stability of the system via Hopf bifurcations. The limit cycle minima and maxima are visualized in green. Stable (unstable) equilibria are marked by solid (dotted) lines. The dashed horizontal line marks the initialization point used for B. **(B)** In the bistable regime, positive and negative stimuli switch the system between sustained bursting and sustained regular firing. Model parameters: Δ = 2, 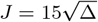, *τ* = 1, *τ*_*A*_ = 10, *α* = 1.0.

## V. DISCUSSION

In this work, we examined the dynamic impact of two different short-term adaptation mechanisms on the collective behavior of a globally coupled QIF population: a multiplicative and an additive one. For both of these mechanisms, we derived and validated meanfield descriptions of the macroscopic dynamics via the approach described in^15^. Using bifurcation analysis, we identified and characterized regimes of collective bursting that emerged given a sufficiently strong adaptation rate. These bursting regimes could co-exist with non-bursting regimes, allowing for dynamic phase transitions between bursting and steady-state behavior via transient inputs. Such bi-stable regimes may be used to describe (1) short-term or working memory as a dynamic property of the collective behavior of neural populations^23^, or (2) transitions between healthy and pathological neurodynamic states such as Epilepsy or Parkinson’s disease^4,5^. Furthermore, due to the boundedness of the bursting regime in 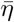, it allows to implement transient forms of population bursting as a response to strong excitatory or inhibitory inputs to the population. Indeed, it has been suggested that bursting in neural populations can be explained by an interaction between alterations in the average population input and post-synaptic homeostatic plasticity^26^. Since the bursting frequency and amplitude scale with the input strength, such types of transient population bursting may even be used to differentiate between different inputs. In summary, this renders our mean-field models applicable to a broad range of neurodynamic scenarios. They provide a description of the emergence of synchronous, bursting neural dynamics in recurrently connected populations of spiking neurons that could either arise from spike-frequency adaptation (additive), or post-synaptic efficacy reduction (multiplicative). Experimental work suggests that the former is a result of different balancing currents which are triggered at a single cell after it generated a spike^25,27^. The latter, on the other hand, has been linked to various mechanisms such as receptor desensitization^16,28^, receptor density reduction^29,30^, or resource depletion at glial cells involved in synaptic transmission^31,32^. Even though these adaptation mechanism can express tremendously different time scales, ranging from a few hundred milliseconds (e.g. spike-frequency adaptation^27^) to days (e.g. post-synaptic receptor density reduction^30^), our meanfield descriptions remain applicable. Furthermore, the results of our bifurcation analysis will hold for different adaptation time scales, as long as *α* is re-scaled accordingly. Finally, our mean-field derivations are independent of the particular form of adaptation that is used. That is, the convolution with an alpha kernel we employed as a second-order approximation of the dynamics of the adaptation variable *A* could be replaced with any other description. This allows future studies the examination of the influence of specific short-term adaptation characteristics on population dynamics.

## ACKNOWLEDGMENTS

The authors thank E. Montbrió for valuable comments and discussions. HS was support by the German Research Foundation (DFG (KN 588/7-1) awarded to TRK, via Priority Program 2041 ‘Computational Connectomics’). Richard Gast has been supported by the Max Planck Society and is currently funded by the Studienstiftung des Deutschen Volkes.

